# Phage infection restores PQS signaling and enhances growth of a *Pseudomonas aeruginosa lasI* quorum-sensing mutant

**DOI:** 10.1101/2021.03.31.438005

**Authors:** Nina Molin Høyland-Kroghsbo, Bonnie L. Bassler

## Abstract

Bacteriophage (phage) therapy is reemerging as a valuable tool to combat multidrug resistant bacteria. A major hurdle in developing efficacious bacteriophage therapies is that bacteria acquire resistance to phage killing. In this context, it is noteworthy that quorum sensing (QS), the bacterial cell-to-cell communication mechanism that promotes collective undertaking of group behaviors including anti-phage defenses, enhances bacterial survival in the face of phage attack. QS relies on the production, release, accumulation, and detection of signal molecules called autoinducers. In the opportunistic pathogen *Pseudomonas aeruginosa*, the LasI/R QS system induces the RhlI/R QS system, and these two systems control, in opposing manners, the PQS QS system that relies on the autoinducer called PQS. A Δ*lasI* mutant is impaired in PQS synthesis, leading to accumulation of the precursor molecule HHQ. HHQ suppresses growth of the *P. aeruginosa* Δ*lasI* strain. We uncover a phage infection-induced mechanism that restores expression of the *pqsH* gene in the *P. aeruginosa* Δ*lasI* QS mutant. PqsH converts HHQ into PQS, preventing HHQ-mediated growth inhibition. Thus, phage-infected *P. aeruginosa* Δ*lasI* cells exhibit superior growth compared to uninfected cells. Phage infection also restores expression of virulence factors and the CRISPR-*cas* anti-phage defense system in the *P. aeruginosa* Δ*lasI* strain. This study highlights a challenge for phage therapy, namely that phage infection may make particular bacterial strains faster growing, more virulent, and resistant to phage killing.

**Importance:** The emergence of multidrug resistant bacteria necessitates development of new antimicrobial therapies. Phage therapy relies on exploiting phages, natural enemies of bacteria, in the fight against pathogenic bacteria. For successful phage therapy development, potent phages that exhibit low propensity for acquisition of bacterial resistance are desired. Here, we show that phage infection restores QS, a cell-to-cell communication mechanism in a *P. aeruginosa* QS mutant, which increases its virulence and resistance to phage killing. Importantly, clinical isolates of *P. aeruginosa* frequently harbor mutations in particular QS genes. Thus, phage therapies against such *P. aeruginosa* strains may inadvertently increase bacterial virulence. Our study underscores the importance of characterizing phage-host interactions in the context of bacterial mutants that are relevant in clinical settings prior to selecting phages for therapy.

## Introduction

Quorum sensing (QS) is a bacterial cell-cell communication process that enables bacteria to collectively control gene expression and orchestrate group behaviors. QS relies on the production and release of signaling molecules called autoinducers (AIs) that accumulate at high cell density and, following detection, activate signal transduction cascades (reviewed in (1)). The opportunistic human pathogen *Pseudomonas aeruginosa* harbors two canonical LuxI/R type QS synthase-receptor pairs, LasI/R and RhlI/R (2, 3). LasI synthesizes the AI 3-oxo-C12-homoserine lactone (3OC12-HSL). 3OC12-HSL interacts with its partner receptor, LasR, and the complex activates expression of genes encoding virulence factors (2, 4, 5). LasR-3OC12-HSL also activates expression of genes encoding the RhlI/R QS system as well as the genes encoding the *Pseudomonas* quinolone signal (PQS) QS pathway (6, 7). RhlI synthesizes the AI C4-homoserine lactone (C4-HSL) that, when bound by RhlR, launches expression of a second set of virulence genes. The RhlR-C4-HSL complex inhibits the PQS QS system (8). PqsA-D are responsible for synthesis of 2-heptyl-4-quinolone (HHQ), which is subsequently converted by PqsH into 2-heptyl-3-hydroxy-4-quinolone, the AI called PQS (9). PQS interacts with its partner receptor, PqsR, and the complex controls downstream gene expression (10).

Bacteriophages (phages) are viruses that infect bacteria. Temperate phages can undergo either lytic development, in which, following infection, they replicate and lyse the host, or lysogenic development, in which the phage integrates into the host genome and becomes a prophage. In response to particular stress cues, prophages can be induced to enter the lytic pathway (reviewed in (11)). Cues governing the lytic pathway generally inform the prophage about the metabolic status and viability of the host bacterium (12–14). Prophages can also “eavesdrop” on host QS signaling and launch their lytic cycles exclusively at high cell density, presumably a condition that optimizes transmission to neighboring bacterial cells (15). Phages harboring QS genes exist in both Gram-positive and Gram-negative bacteria (16, 17). Additionally, phages can communicate with one another to modulate their lysis-lysogeny transitions (18, 19).

We previously reported that *P. aeruginosa* QS, through LasI and RhlI, increases expression, activity, and adaptation capability of the clustered regularly interspaced short palindromic repeats (CRISPR) and the CRISPR associated (Cas) phage defense components in *P. aeruginosa* UCBPP-PA14 (called PA14) (20). We speculated that this regulatory mechanism ensures maximal expression of phage defenses at high cell density, when the risk of phage infection is high due to the proximity of many host bacteria. Phage receptors are often QS regulated, which can confound interpretation of phage-QS interactions. To circumvent this issue, in our previous study, we measured the ability of CRISPR-Cas to eliminate a foreign plasmid (20).

Depending on whether the infecting agent is a plasmid or a phage, the mechanisms underlying parasitism and the outcomes to the bacterial prey differ. Here, we extend our analyses to the effects of QS on phage infection. We focus on infection of WT PA14 and a PA14 Δ*lasI* QS mutant by the phage JBD44, which employs the O-antigen as its receptor, as importantly, the O-antigen is not QS regulated (21, 22). To our surprise, phage JBD44 killed WT PA14 more efficiently than the QS mutant. Specifically, after an initial killing phase, phage JBD44 infection enhanced the growth of the PA14 Δ*lasI* strain. Restoration of expression of *lasR* and genes in the Rhl and PQS QS pathways that function downstream of the LasI/R QS system drove the enhanced bacterial growth. Moreover, phage infection restored expression of QS-activated genes encoding virulence factors and the genes in the CRISPR-*cas* immune defense system.

## Results

### Phage infection enhances growth of the *P. aeruginosa* Δ*lasI* QS mutant

Our goal was to investigate if and how phage infection and QS, together, affect PA14 phenotypes. For such an analysis, a phage that does not use a QS-regulated receptor for infection was required. Thus, we exploited phage JBD44 that uses the non-QS-regulated O-antigen as its receptor (21, 22) so it exhibits equal infectivity irrespective of host cell QS status.

We assessed the plaque morphology of phage JBD44-infected PA14. JBD44 is a temperate phage so it can enact the lytic program or it can integrate into the host chromosome. The latter process generates lysogenic cells, which are immune to killing by the same or closely related phages. Consequently, infection by phage JBD44 results in turbid plaques on lawns of PA14 (Fig. 1A). By contrast, infection by virulent phages that cannot form lysogens results in clear plaques. Virulent phages are desirable for phage therapies because they are obligate killers. With possible applications in mind, we isolated a spontaneous small-clear-plaque mutant of phage JBD44, which could be a candidate virulent phage (Fig. 1B). Sequencing revealed that the mutant phage possesses an Asn to Lys alteration at residue 289 (N289K) in Gp39. We call the isolated mutant phage JBD44^39*^. Gp39 is a putative tail fiber protein. Some phage tails possess depolymerase activity against bacterial capsules and this capability drives halo formation around plaques (23). Indeed, WT phage JBD44 makes plaques with halos and the mutant JBD44^39*^ phage does not (Fig. 1A and 1B, respectively). We suspect that the WT phage JBD44 phenotype could be due to Gp39 LPS depolymerase activity causing capsule degradation. Propagation of phage JBD44^39*^ on PA14 led to rare spontaneous revertants of JBD44^39*^ that restored the WT large turbid plaque morphology including the surrounding halos. We sequenced one phage revertant. It encoded a Gp39 K289T substitution, suggesting that position 289 in GP39 is critical for Gp39 to drive plaque size, turbidity, and halo formation.

**Fig. 1.**
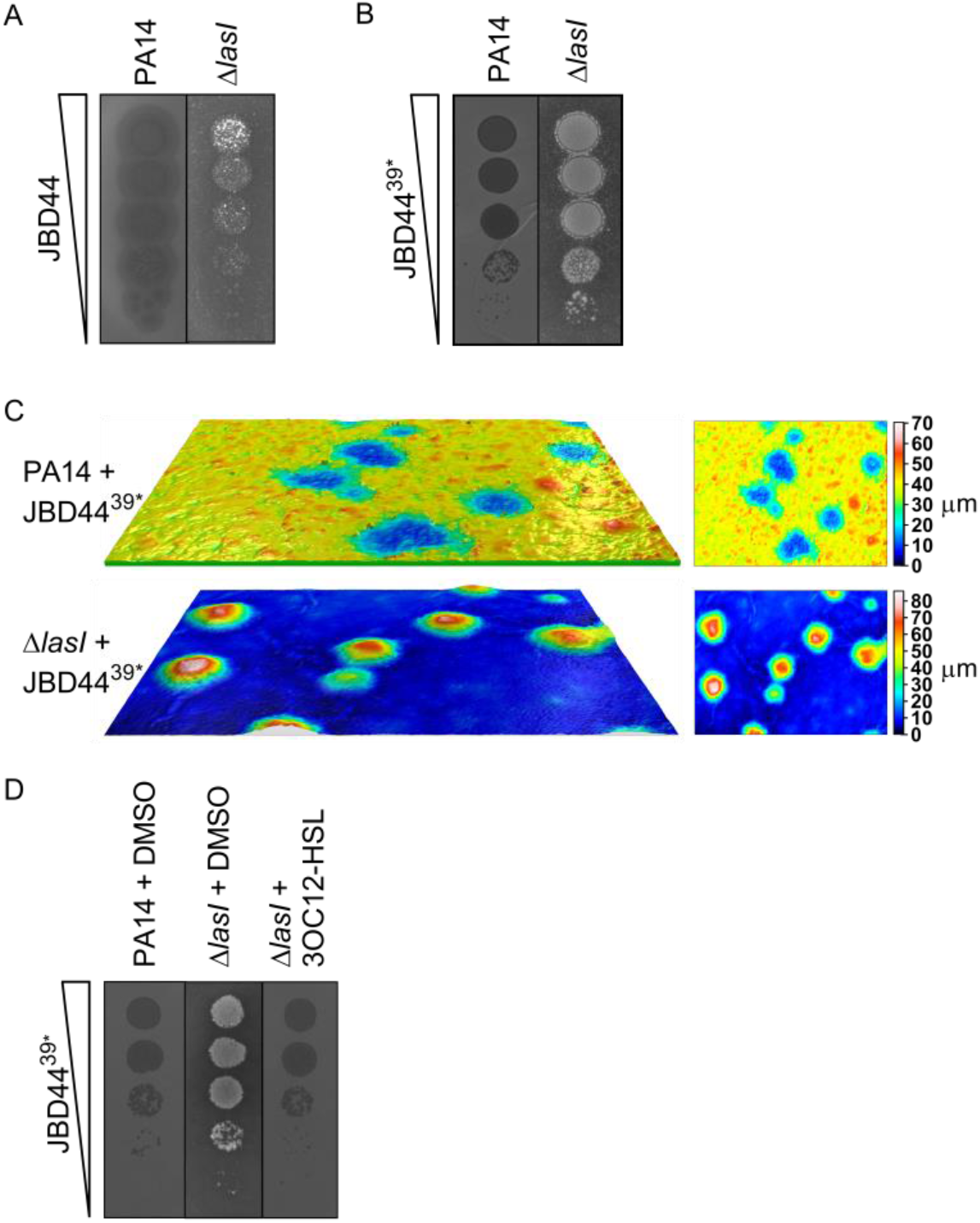
Phage infection causes reverse plaque formation on the PA14 Δ*lasI* mutant. Plaque assay showing 10-fold serial dilutions of phages JBD44 (A) and JBD44^39*^ (B) spotted onto lawns of PA14 and the PA14 Δ*lasI* mutant. (C) Surface profiles of individual plaques from panel B, measured using a Leica DCM 3D Micro-optical System. The left images show side views of plaques on lawns of PA14 (top) and the PA14 Δ*lasI* mutant (bottom). The right images show top views of the same plaques. Scale bars show the heights (μm) of the lawns and plaques with black/blue representing the lowest points and red/white representing the highest points. (D) Plaque assay showing 10-fold serial dilutions of phage JBD44^39*^ spotted onto lawns of PA14 and the PA14 Δ*lasI* mutant treated with DMSO (control solvent) or 2 μM 3OC12-HSL.

QS regulates multiple phage defenses (20, 24–27). Thus, we investigated whether the JBD44 and JBD44^39*^ phages drove identical plaquing efficiencies and plaque morphologies on PA14 and on a QS mutant. To explore this question, we chose the PA14 Δ*lasI* QS mutant because the LasI synthase functions at the top of the QS hierarchy and, thus, the LasI-synthesized 3OC12-HSL AI launches the RhlI/R and PQS QS cascades. As described above, both phages formed plaques on lawns of PA14 (Fig. 1A and 1B). By contrast, on lawns of the PA14 Δ*lasI* strain, both phages promoted bacterial growth (Fig. 1A and 1B). We call the plaque phenotype on the PA14 Δ*lasI* strain the “reverse plaque”. Phage JBD44^39*^-driven reverse plaques achieved a height of approximately 70 μm above the PA14 Δ*lasI* cell lawn (Fig. 1C). By contrast, infection of PA14 with phage JBD44^39*^ resulted in plaques with ~35 μm depressions below the surface of the lawn (Fig. 1C). Normal plaque morphology was restored to the phage JBD44^39*^-infected PA14 Δ*lasI* mutant when exogenous 2 μM 3OC12-HSL was supplied (Fig. 1D). Thus, the absence of the LasI QS pathway is required for development of the reverse plaque phenotype.

### Phage infection of the PA14 Δ*lasI* strain proceeds with an initial phase of lysis followed by reverse plaque formation

To investigate the development of the reverse plaque phenotype on the PA14 Δ*lasI* strain, we continued using phage JBD44^39*^ because it induced a stronger growth enhancement than WT phage JBD44 (Fig. 1A and 1B). First, regarding timing: we imaged JBD44^39*^ plaque development on lawns of PA14 and the PA14 Δ*lasI* strain every 1 h (Movie S1). Fig. 2A shows that infection of PA14 caused low-level lysis of the bacterial lawn at 6 h, followed by maximal lysis by 18 h, and no apparent change in plaque phenotype occurred between 18 and 36 h. Similar results were obtained for the PA14 Δ*lasI* strain at initial times. At the 18 h time point, however, the plaques on the PA14 Δ*lasI* strain began to show turbidity, indicating bacterial growth. Growth inside the plaques continued through 30-36 h, by which time, the reverse plaques had fully formed (Fig. 2A). These data suggest that there are two phases of phage JBD44^39*^ infection of the PA14 Δ*lasI* strain: bacterial lysis at early times followed by enhanced growth of infected host cells. Notably, the PA14 lawn was green due to production of the QS-regulated virulence factor pyocyanin, whereas the lawn of uninfected PA14 Δ*lasI* cells and the lawn surrounding the phage JBD44^39*^-infected PA14 Δ*lasI* strain lacked the green hue due to the absence of pyocyanin production (Fig. 2A and 2B). These results were expected since LasI/R is required to activate the RhlI/R QS system which, in turn, induces pyocyanin synthesis (3, 5, 28). Surprisingly, the surviving cells in the reverse plaques on the PA14 Δ*lasI* strain produced pyocyanin (Fig. 2A and 2B). Likewise, restoration of pyocyanin production occurred when PA14 Δ*lasI* cells were supplied 3OC12-HSL (Fig. 2B). Together, these results suggest that phage JBD44^39*^ infection of the PA14 Δ*lasI* strain reactivates the QS pathway.

**Fig. 2.**
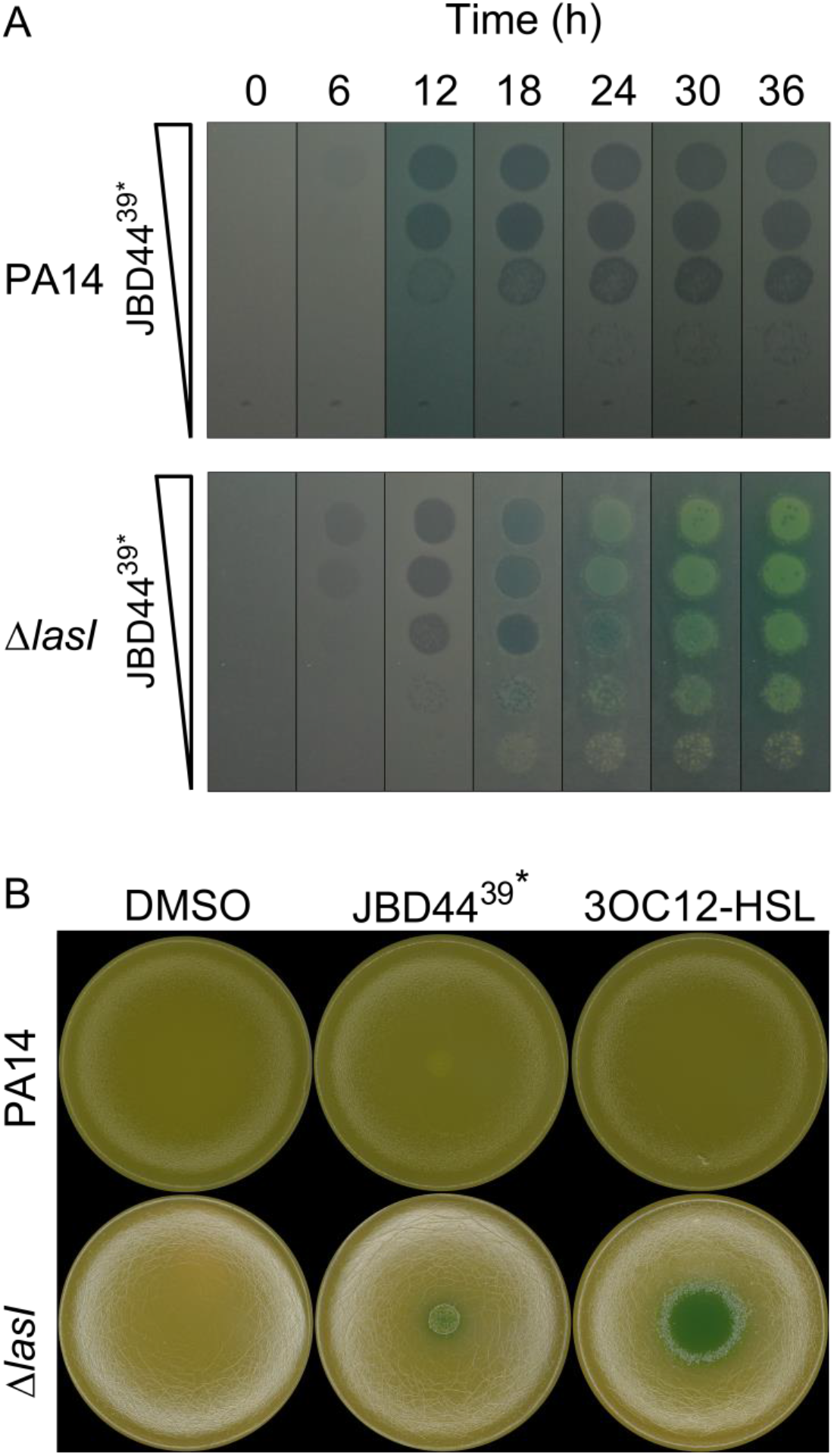
Time series and pyocyanin production profiles following phage JBD44^39*^-driven plaque development on PA14 and the PA14 Δ*lasI* strain. (A) Plaque assay showing 10-fold serial dilutions of phage JBD44^39*^ spotted onto lawns of PA14 (top panel) and the PA14 Δ*lasI* strain (bottom panel). These data are also shown in Supplementary Movie S1. (B) Lawns of PA14 (top row) and the PA14 Δ*lasI* strain (bottom row) to which 5 μL DMSO, JBD44^39*^, or 100 μM 3OC12-HSL was added at the center of the plate. Pyocyanin appears as a blue-green colored pigment.

### The PA14 Δ*lasI* cells in the reverse plaques are phage JBD44^39*^ lysogens

Reverse plaque formation on the PA14 Δ*lasI* strain suggested that phage JBD44^39*^ is not virulent to the QS mutant. One possibility is that phage JBD44^39*^ lysogenizes the PA14 Δ*lasI* strain. To test this notion, we isolated six single colonies from the reverse plaques. These isolates were all resistant to phage killing by phage JBD44^39*^, as judged by the inability of phage JBD44^39*^ to form plaques on them (Fig. 3A). Moreover, all six isolates released JBD44^39*^ phage particles in liquid culture as measured by the ability of the chloroform-sterilized conditioned media prepared from the isolates to form plaques on lawns of PA14 and to form reverse plaques on lawns of the PA14 Δ*lasI* strain (Fig. 3B). Therefore, the PA14 Δ*lasI* cells in the reverse plaques are phage JBD44^gp39*^ lysogens. Interestingly, the colony morphology of PA14 is smooth, whereas that of the PA14 Δ*lasI* strain and the PA14 Δ*lasI* JBD44^39*^ lysogens are rough with a metallic, iridescent sheen (Fig. 3C). This phenotype resembles the autolytic phenotype reported for *P. aeruginosa lasR* mutants (29, 30). We return to this point below.

**Fig. 3.**
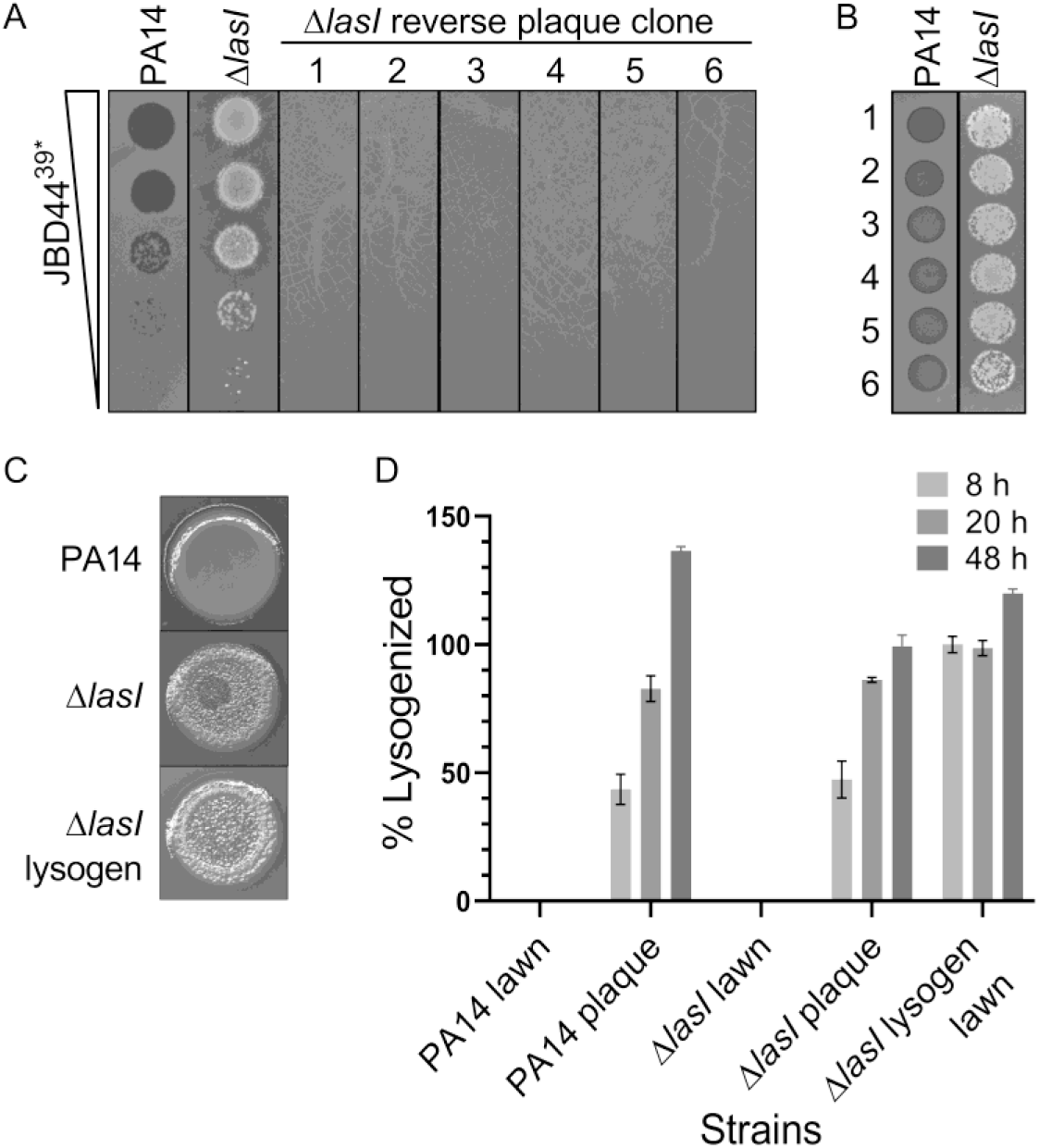
Phage JBD44^39*^ lysogenizes both PA14 and the PA14 Δ*lasI* strain. (A) Plaque assay showing 10-fold serial dilutions of phage JBD44^39*^ on lawns of PA14, the PA14 Δ*lasI* strain, and six isolates from reverse plaques from the PA14 Δ*lasI* strain that had been infected with phage JBD44^39*^. (B) Cell-free fluids prepared from overnight cultures of the six isolates shown in panel A spotted onto lawns of PA14 and the PA14 Δ*lasI* strain. (C) Colony morphologies of PA14, the PA14 Δ*lasI* strain, and the PA14 Δ*lasI* strain lysogenized with phage JBD44^39*^ grown from 5 μL overnight cultures (each image is 1 cm wide). (D) The percentage of cells lysogenized by phage JBD44^39*^ at 8, 20, and 48 h. The first set of bars represents uninfected lawns of PA14, the second set represents JBD44^39*^ plaques on PA14, the third set shows uninfected lawns of the PA14 Δ*lasI* strain, the fourth set represents JBD44^39*^ reverse plaques on the PA14 Δ*lasI* strain, and the fifth set is from lawns of the PA14 Δ*lasI* strain that was already lysogenized by phage JBD44^39*^. The percentage of lysogenized cells was measured by qPCR of genomic DNA using primers specific for the phage JBD44 integration site. Data are normalized to the reference gene encoding the 5S ribosomal subunit. Error bars designate standard deviations from *n* = 3 biological replicates.

To monitor phage JBD44^39*^-lysogenization of cells over time, we sampled the centers of plaques following phage JBD44^39*^ infection of PA14 and the PA14 Δ*lasI* strain at 8, 20, and 48 h. We likewise sampled regions of the surrounding uninfected lawns of PA14, the PA14 Δ*lasI* strain, and the PA14 Δ*lasI* phage JBD44^39*^ lysogen. We measured lysogenization using qPCR amplification of the phage JBD44 integration site in the PA14 chromosome. Not surprisingly, the uninfected PA14 and PA14 Δ*lasI* lawns showed no lysogenization by phage JBD44^39*^, and the PA14 Δ*lasI* JBD44^39*^ lysogen was fully lysogenized (Fig. 3D). Regarding the samples from inside the plaques, the percentage of lysogenized cells rose similarly for both strains over time (Fig. 3D). Roughly half of each population was lysogenized by 8 h and all of each population was lysogenized by 48 h (Fig. 3D). This result demonstrates that while JBD44^39*^ forms clear plaques on PA14, it is not virulent, and is indeed capable of lysogenizing the cells. Therefore, the differences in plaque morphologies between the two strains (Fig. 3A) are not due to differences in lysogenization frequency, but rather, may be caused by differences in the strains’ responses to phage infection and/or differences in the strains' responses to lysogenization.

### CRISPR-*cas* is not required for autolysis or JBD44^39*^-induced reverse plaque formation on the PA14 Δ*lasI* strain

We wondered if the autolytic trait of the PA14 Δ*lasI* strain and the ability of phage JBD44^39*^ to form reverse plaques were connected to the CRISPR-Cas system. Autolysis can be induced by activation of prophages residing within bacterial host genomes. Indeed, PA14 lysogenized by phage DMS3 autolyses when grown as a biofilm, due to autoimmunity stemming from CRISPR-Cas directed targeting of the DMS3 prophage in the host chromosome (31). Phage JBD44 is not targeted by the PA14 CRISPR-Cas system because the PA14 CRISPR arrays do not encode CRISPR RNAs targeting this phage. However, CRISPR adaptation against phage JBD44^39*^could occur. If so, adaptation would drive subsequent targeting of phage JBD44^39*^. To test whether CRISPR-Cas is required for autolysis and for phage JBD44^39*^ to drive reverse plaque formation on the PA14 Δ*lasI* strain, we infected a PA14 Δ*lasI* ΔCRISPR Δ*cas* mutant with phage JBD44^39*^. Fig. 4 shows that there is no difference in plaque morphology between the PA14 Δ*lasI* strain and the PA14 Δ*lasI* ΔCRISPR Δ*cas* strain following infection, and like the PA14 Δ*lasI* strain, the PA14 Δ*lasI* ΔCRISPR Δ*cas* strain is autolytic. These results eliminate any role for CRISPR-Cas in in reverse plaque formation and autolysis on this strain.

**Fig. 4.**
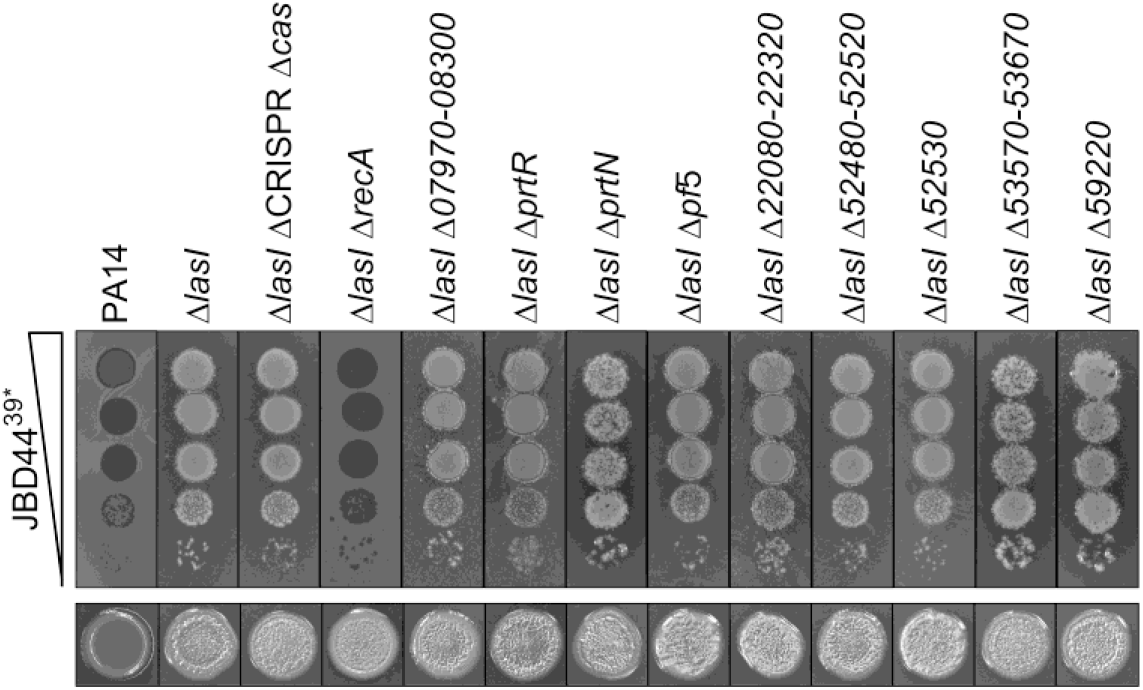
RecA is required for phage JBD44^39*^-directed reverse plaque formation on the PA14 Δ*lasI* mutant but not for autolysis. Top panel: Plaque assays showing 10-fold serial dilutions of phage JBD44^39*^ spotted onto lawns of PA14, the PA14 Δ*lasI* strain, and the PA14 Δ*lasI* strain with the designated gene(s) also deleted. Bottom panel: Colony morphologies of the designated strains grown from 5 μL overnight cultures (each image is 1 cm wide).

### RecA is required for phage JBD44^39*^-induced reverse plaque formation on the PA14 Δ*lasI* strain but not for the autolysis phenotype

Prophages can provide the bacteria within which they reside resistance to infection by related phages, a phenomenon called superinfection exclusion (32). In the case of *Pseudomonas*, broad superinfection exclusion can occur due to the combined effects of the expression of prophage genes encoding diverse defenses against incoming phages (21). We wondered whether prophage genes specifying superinfection exclusion factors or other defense components are activated in the PA14 Δ*lasI* strain but not in PA14, and, if so, if those genes could underpin the autolysis and reverse plaque formation phenotypes. RecA is required for induction of prophage genes. Thus, we deleted *recA* from the PA14 Δ*lasI* strain and subsequently, infected it with phage JBD44^39*^. In the absence of *recA*, phage JBD44^39*^ formed small clear plaques, demonstrating that RecA is indeed required for reverse plaque formation and, moreover, suggesting a prophage could be involved in driving these phenotypes (Fig. 4, top panel). The Δ*lasI* Δ*recA* mutant was autolytic, so RecA is not required for that process (Fig. 4, bottom panel).

To test the possibility that RecA-activated prophage components promote reverse plaque formation, we individually eliminated all prophages and prophage elements from the PA14 Δ*lasI* strain and assayed colony and plaque morphologies following infection by phage JBD44^39*^. Specifically, we deleted the prophage *PA14_07970-08300* and the genes *prtR* and *prtN* encoding its repressor and activator, respectively, prophage Pf5 (*PA14_48880-49030*) (33), the cryptic prophage element *PA14_22080-22320,* the cryptic prophage operon *PA14_52480-52520,* the gene encoding its repressor *PA14_52530*, the cryptic prophage element *PA14_53570-53670*, and the S5 Pyocin (*PA14_59220*). Reverse plaques formed on all of the deletion mutants and all the mutants remained autolytic. These results show that, individually, none of prophages or prophage components drives reverse plaque development or autolysis (Fig. 4). Thus, RecA is dispensable for autolysis but key for reverse plaque formation in the PA14 Δ*lasI* strain, but at present, for unknown reasons.

### PqsA is required for phage JBD44^39*^reverse plaque formation on the PA14 Δ*lasI* strain

We hypothesize that phage JBD44^39*^ infection activates QS in the PA14 Δ*lasI* strain because pyocyanin production was restored (Fig. 2B). To explore this notion, and moreover, to define which QS components are involved, we assayed the plaque morphologies of phage JBD44^39*^-infected PA14 deletion mutants in the Las, Rhl, and PQS QS pathways as well as in the gene encoding the orphan QS receptor QscR (34). As in Fig. 1, phage JBD44^39*^ formed clear plaques on lawns of PA14 and reverse plaques on the PA14 Δ*lasI* strain (Fig. 5). Reverse plaque formation was less pronounced on the PA14 strains lacking Δ*lasR* than on the PA14 Δ*lasI* strain (Fig. 5, top panel). Usually, a mutant lacking a QS synthase has the identical phenotype as the mutant lacking the cognate receptor, because AIs and partner receptors function as obligate pairs (reviewed in (1)). However, there are exceptions in *P. aeruginosa*, potentially explaining the differences in plaque morphologies between the PA14 Δ*lasI* and the PA14 Δ*lasR* strains (20, 35, 36). Reverse plaques formed on all strains lacking *lasI*, except for the PA14 Δ*lasI* Δ*pqsA* strain, in which phage JBD44^39*^ plaques were small and clear, similar to those formed on PA14 (Fig. 5, top panel). Thus, PqsA is critical for reverse plaques to form on the PA14 Δ*lasI* mutant. In every QS mutant strain we tested, the ability of phage JBD44^39*^ to cause reverse plaque formation always correlated with the autolytic phenotype, exhibited by rough and iridescent colony surfaces (Fig. 5, bottom panel) suggesting that the iridescent surface precipitate could be required for reverse plaque formation.

**Fig. 5.**
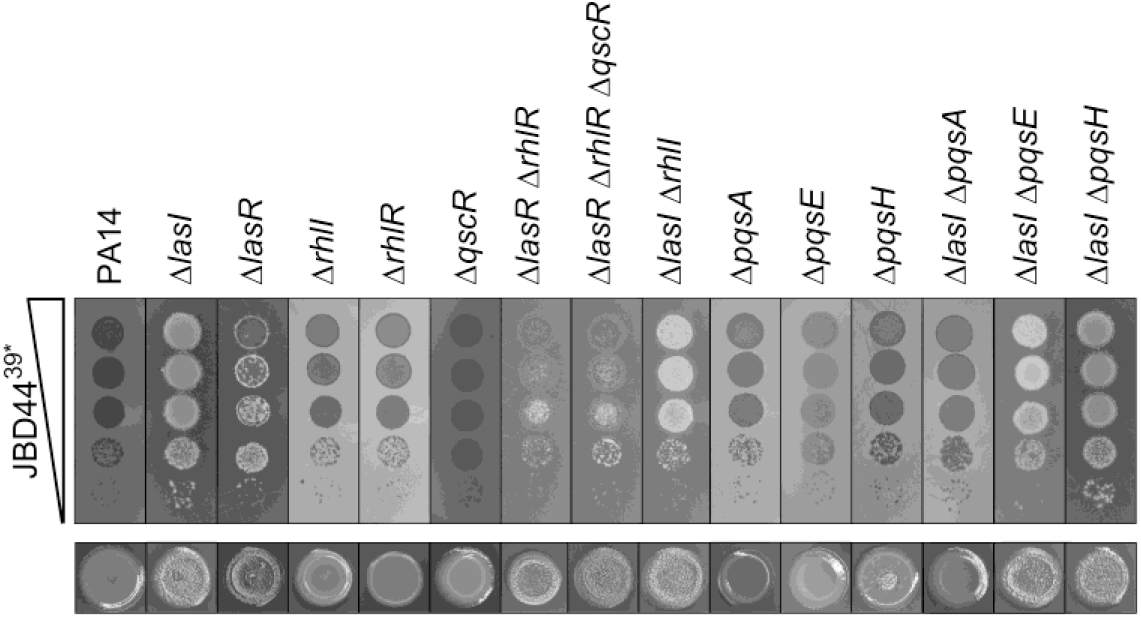
PqsA is required for phage JBD44^39*^-directed autolysis and reverse plaque formation on the PA14 Δ*lasI* strain. Top panel: Ten-fold serial dilutions of phage JBD44^39*^ were spotted onto lawns of PA14 and the designated mutants. Bottom panel: Colony morphologies of the designated strains grown from 5 μL overnight cultures (each image is 1 cm wide).

### Phage JBD44^39*^ infection restores transcription of QS pathway genes and PQS production in the PA14 Δ*lasI* mutant

The requirement for *pqsA* in phage JBD44^39*^-driven autolysis and reverse plaque formation on the PA14 Δ*lasI* strain, coupled with our finding that reverse plaques produce pyocyanin, which is normally made in response to LasI/R-mediated activation of PQS QS (Fig. 2B), suggested that phage JBD44^39*^ infection could activate the PQS QS system in the absence of LasI. To explore this possibility, we tested whether phage JBD44^39*^ infection of the PA14 Δ*lasI* strain affected expression of QS genes. RT-qPCR showed that, as expected given what is known about the arrangement of QS regulatory components, the PA14 Δ*lasI* strain exhibited reduced expression of the QS pathway components *lasR*, *rhlR*, *rhlI*, *pqsH*, and *pqsA* compared to PA14 (Fig. 6A). Phage JBD44^39*^ infection caused little change in expression of QS genes in PA14. By contrast, phage JBD44^39*^ infection of the PA14 Δ*lasI* strain restored expression of all of the above QS genes to at least 50% of their WT levels (Fig. 6A). We highlight that *pqsA* expression in the phage-infected PA14 Δ*lasI* strain increased ~7-fold above the WT level. We discuss a potential mechanism underlying this result in the next paragraph. As noted earlier, PqsA-D synthesize the PQS precursor molecule HHQ, which, in a final biosynthetic step, is converted into PQS by PqsH. With respect to *pqsH*, in the phage JBD44^39*^-infected PA14 Δ*lasI* mutant, expression changed from undetectable to 50% of the PA14 level, potentially restoring the ability of the bacterium to convert HHQ into PQS.

**Fig. 6.**
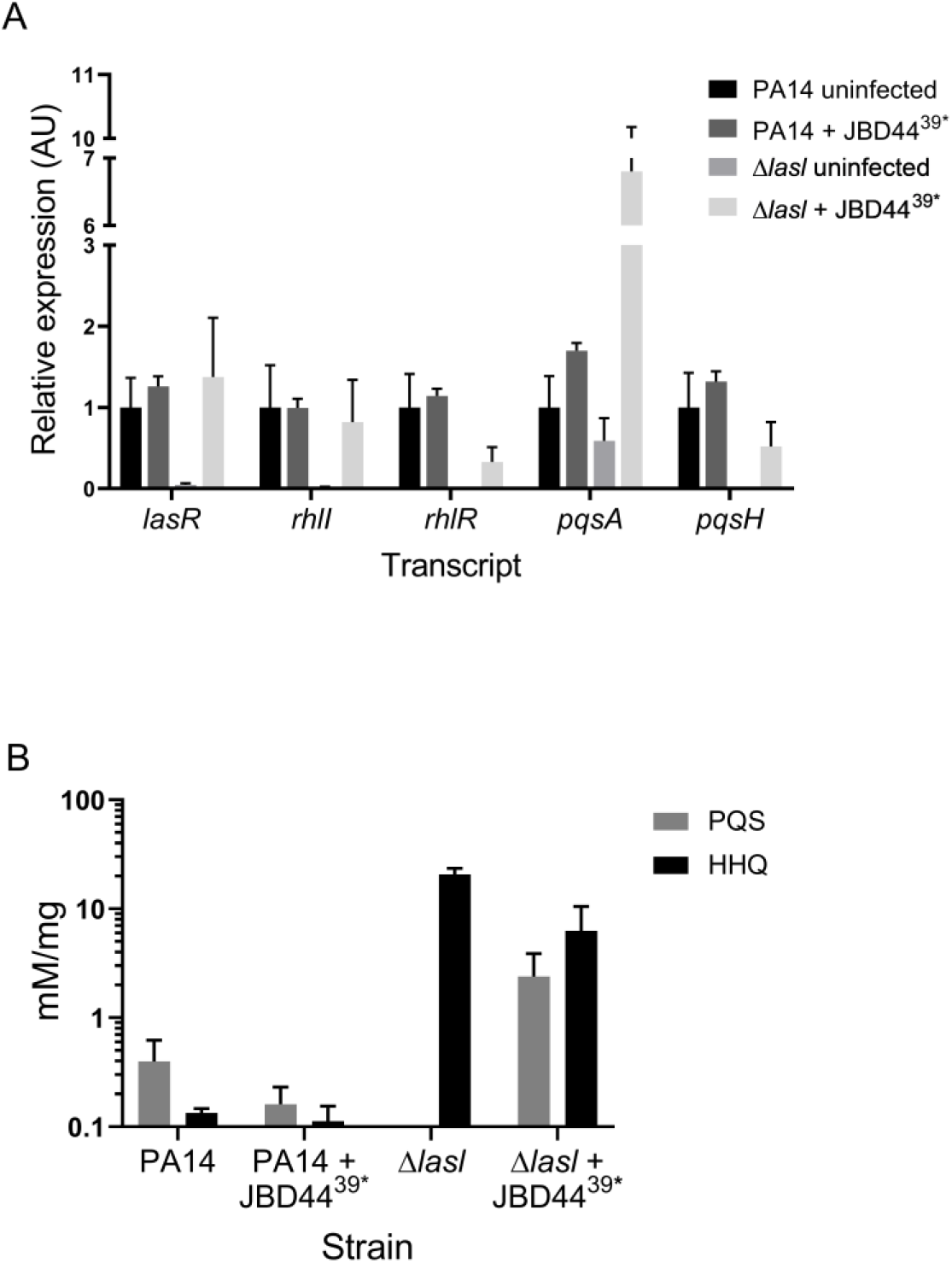
Phage JBD44^39*^-infection activates expression of QS pathway genes and restores PQS production to the PA14 Δ*lasI* mutant. (A) Shown are transcript levels of the designated genes in uninfected- and phage JBD44^39*^-infected lawns of PA14 and the PA14 Δ*lasI* strain. Relative transcript levels were normalized to 5S RNA. Error bars designate standard deviations from *n =* 3 biological replicates. (B) Relative levels of HHQ and PQS present in cells from uninfected lawns and from plaques formed by phage JBD44^39*^-infected PA14 and the PA14 Δ*lasI* strain. PQS and HHQ levels were measured by liquid chromatography-mass spectrometry per mg lawn or per mg plaque material and the relative concentrations were quantified using commercial PQS and HHQ standards. One cannot compare levels across the conditions, only within each condition (uninfected lawns or plaques). Error bars represent standard deviations from *n* = 3 biological replicates.

Our finding of potential differences in HHQ production in the phage-infected mutants under study made us wonder if the molecule responsible for the iridescent sheen on the PA14 Δ*lasI* strain could be HHQ, and if so, if HHQ could drive autolysis. Evidence for this possibility comes from earlier studies showing that *P. aeruginosa lasR* null mutants overproduce HHQ compared to WT (29, 30). Indeed, the PA14 Δ*lasR* strain exhibited both the sheen and the autolytic phenotype similar to the PA14 Δ*lasI* strain (Fig. 5, bottom panel). We next considered whether upregulation of *pqsH* expression in the JBD44^39*^-infected PA14 Δ*lasI* strain (Fig. 6A) resulted in PqsH-mediated conversion of HHQ into PQS. To investigate this possibility, we measured the relative levels of HHQ and PQS in uninfected lawns of PA14 and the PA14 Δ*lasI* strain and in phage JBD44^39*^-infected plaques of PA14 and the PA14 Δ*lasI* strain using liquid chromatography-mass spectrometry. Because the uninfected and infected strains grew to different levels, direct comparisons of compound per weight of material could not be made. To circumvent this issue, we compared the ratios of PQS to HHQ in each strain under each condition. Lawns of PA14 cells possessed 3-fold more PQS than HHQ, irrespective of phage infection state. By contrast, lawns of the PA14 Δ*lasI* strain possessed HHQ but had no detectable PQS (Fig. 6B), similar to previous observations (9, 30). Importantly, reverse plaques of the JBD44^39*^-infected PA14 Δ*lasI* strain possessed readily detectable PQS, although nearly 3-fold more HHQ than PQS was present (Fig. 6B). Thus, phage JBD44^39*^ infection of the PA14 Δ*lasI* strain restored the ability of the strain to convert HHQ into PQS. PQS binding to PqsR launches a positive feedback loop that activates expression of *pqsABCDE*. We expect that this feedback loop underlies the high expression of *pqsA* that occurs in the JBD44^39*^-infected PA14 Δ*lasI* strain (Fig. 6A). Moreover, these results explain why deletion of *pqsA* in the PA14 Δ*lasI* strain abrogated autolysis (Fig. 5, bottom panel): this mutant makes no HHQ. Our data are also in agreement with the observation that a *P. aeruginosa* PAO1 *lasR* mutant produces PQS in late stationary phase growth (37). Thus, under particular physiological conditions, including late stationary phase growth and following phage infection, the PQS QS pathway can be activated independently of LasI/R QS signaling.

### Phage JBD44^39*^ infection restores expression of virulence genes and *cas3* in the PA14 Δ*lasI* strain

QS governs expression of genes that collectively benefit bacterial communities, e.g. virulence genes and phage defense genes such as CRISPR-*cas*. Given that transcript levels of QS regulatory genes increased following phage JBD44^39*^ infection of the PA14 Δ*lasI* strain (Fig. 6A), it follows that expression of QS-regulated target genes could also be upregulated. To examine this possibility, we assayed *lasB*, *rhlA*, and *phzA* expression as readouts of QS-directed virulence activity. The LasI/R QS system promotes expression of *lasB,* encoding the extracellular virulence factor elastase (38, 39). The rhamnolipid biosynthesis gene *rhlA* is activated by the Rhl QS system (39). PQS QS upregulates expression of the *phzA* gene involved in production of phenazines including pyocyanin, which plays a role in biofilm development and virulence (40, 41).

As expected, all three QS-regulated virulence genes were downregulated in the PA14 Δ*lasI* mutant, with 3,000-fold reduced *lasB*, 60-fold reduced *rhlA*, and 7-fold reduced *phzA* expression compared to PA14 (Fig. 7). In the phage JBD44^39*^-infected PA14 Δ*lasI* strain, *lasB* expression increased to 20% of the level in PA14 and *rhlA* expression was fully restored. *phzA* was also upregulated 3-fold and 84-fold following phage JBD44^39*^ infection of PA14 and the PA14 Δ*lasI* strain, respectively, suggesting that phage infection can potentially enhance virulence of PA14. Phage JBD44^39*^-infection mediated induction of expression of *phzA* in the PA14 Δ*lasI* strain exceeded that in the infected PA14 strain. *phzA* expression is activated by PQS, so this finding is consistent with the observed heightened expression of *pqsA* and restoration of PQS production in the JBD44^39*^-infected Δ*lasI* strain (Fig. 6A and B). *phzA* encodes a pyocyanin biosynthesis pathway component, also in agreement with our observation of pyocyanin production in the reverse plaques in the JBD44^39*^-infected Δ*lasI* strain (Fig. 2B). Moreover, phage JBD44^39*^ infection of PA14 and the PA14 Δ*lasI* strain activated transcription of *cas3* 2-fold and 15-fold, respectively (Fig. 7). These results, are to our knowledge, the first evidence showing a phage infection-mediated increase in virulence and anti-phage defense gene expression in PA14.

**Fig. 7.**
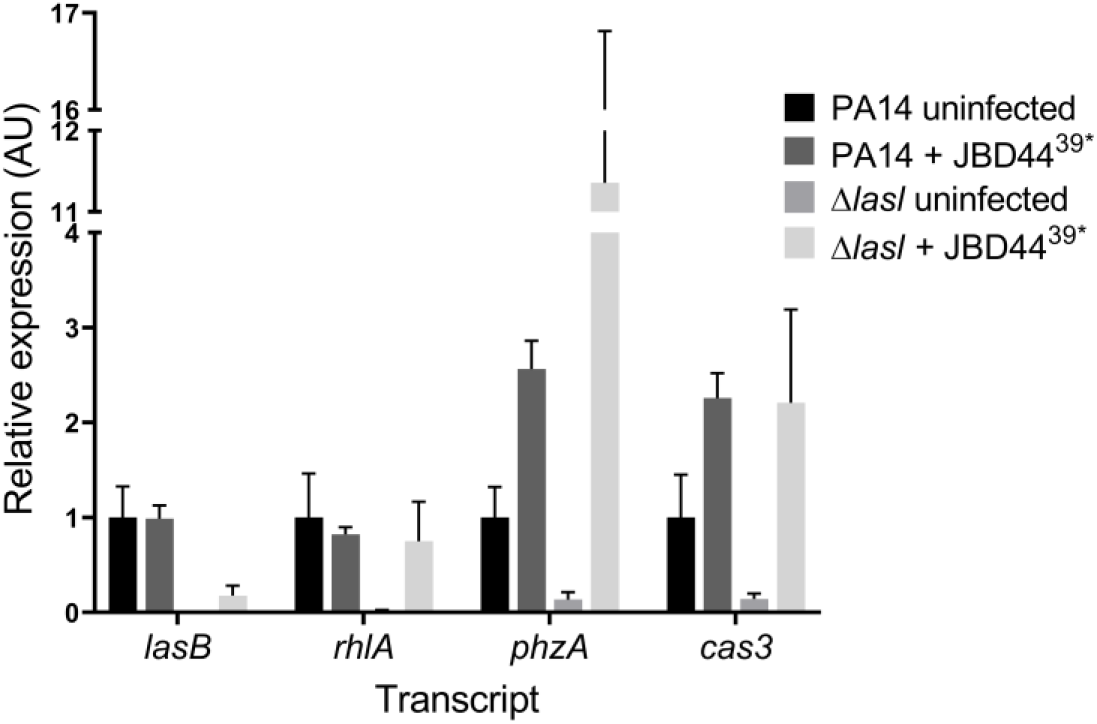
Phage JBD44^39*^ infection activates expression of virulence genes and the anti-phage defense gene *cas3* in the PA14 Δ*lasI* mutant. Shown are transcript levels of the designated genes in uninfected and phage JBD44^39*^-infected PA14 and the PA14 Δ*lasI* strain. Relative transcript levels were normalized to 5S RNA. Error bars designate standard deviations from *n* = 3 biological replicates.

## Discussion

Here, we show that phage JBD44^39*^ infection can increase expression of *lasR* and genes encoding components of the RhlI/R and PQS QS systems, QS regulated virulence genes, and *cas3* in the *P. aeruginosa* PA14 Δ*lasI* strain (Fig. 6 and 7). Phage infection of the PA14 Δ*lasI* strain caused an initial phase of killing, followed by growth enhancement, exceeding that of uninfected surrounding PA14 Δ*lasI* cells, giving rise to reverse plaques (Fig. 1C and 2A). RecA was required for reverse plaque formation (Fig. 4, top panel) but not for autolysis (Fig. 4, bottom panel). Endogenous prophage and phage-derived pyocin genes were, individually, dispensable for both phenotypes, indicating that other genes downstream of RecA may be involved. We found that PqsA is required for both reverse plaque formation and the autolytic phenotype exhibited by the PA14 Δ*lasI* strain (Fig. 5, top panel), and the accumulation of the PqsA-D product, the HHQ molecule (Fig. 5, bottom panel). Phage-mediated upregulation of *pqsH* expression in the PA14 Δ*lasI* strain (Fig. 6A) enabled conversion of HHQ to PQS (Fig. 6B) suggesting requirements for HHQ for autolysis and PQS for reverse plaque formation.

We hypothesize that during initial phage JBD44^39*^-infection of the PA14 Δ*lasI* strain, when the QS pathways are not yet reactivated, phage infection causes normal plaque development. Over time, the PA14 Δ*lasI* cells accumulate HHQ, causing autolysis. We imagine that RecA detects the lysogenization process and/or stress responses induced by phage replication, and by an unknown mechanism, RecA drives increased *pqsH* expression. Consequently, phage JBD44^39*^-infected PA14 Δ*lasI* cells convert HHQ into PQS, which alleviates HHQ toxicity and endows infected cells with superior growth capabilities relative to the surrounding uninfected cells, which continue to suffer from HHQ accumulation and, thus, autolysis.

A crucial finding of ours is that the PA14 Δ*lasI* JBD44^39*^ lysogens exhibit the identical autolytic colony morphology as the PA14 Δ*lasI* mutant (Fig. 3C), suggesting that when the phage exists as a lysogen it does not drive upregulation of *pqsH* and restoration of PQS QS signaling. Rather, active infection of non-lysogenic PA14 Δ*lasI* cells, again, likely detected by RecA, triggers *pqsH* expression, enabling the cells to produce and release PQS, which, in turn, induces *pqsH* expression in neighboring non-infected or lysogenic cells. As a consequence of JBD44^39*^-mediated upregulation of the QS pathways in the PA14 Δ*lasI* strain, QS-dependent activation of expression of *lasB*, *rhlA*, and *phzA* occurred (Fig. 7). Increased *phzA* expression drove increased pyocyanin production (Fig. 2B). JBD44^39*^ infection likewise enhanced expression of *cas3*, encoding the nuclease of the type I-F CRISPR-Cas system (Fig. 7). Thus, JBD44^39*^ infection increases both the virulence potential and phage defense capacity of the PA14 Δ*lasI* strain. Of note, we also observed that reverse plaques form on PA14 Δ*lasI* cells in response to infection by phages DMS3, JBD13, JBD18, and JBD25 (Fig. S1), suggesting that reverse plaque formation, enhanced QS signaling, and expression of QS-regulated traits may be general responses to phage infection in the PA14 Δ*lasI* strain.

Other recent findings also connect bacterial QS to interactions with phages. Bacteria use the accumulation of QS AIs as indicators of impending phage infection, and in response, they modulate their phage defenses to appropriately combat threats (20, 24–27). Phage infection of *P. aeruginosa* PAO1 activates expression of PQS biosynthesis genes (42) and phage infection and antibiotic-induced stress activate PA14 PQS production which allows PQS to function as an alarm signal that alerts nearby uninfected PA14 cells to physically avoid phage-infected or antibiotic-stressed cells (43). In contrast, phage infection of *Enterococcus faecalis* inhibits expression of its Fsr QS system (44). Potentially, the phage used in this *E. faecalis* study harbors an anti-QS gene, similar to a small protein LasR inhibitor recently discovered in the *P. aeruginosa* DMS3 phage (45). Moreover, phages surveil bacterial QS AIs and tune their lysis-lysogeny decisions to host cell density (15). Here, we speculate that HHQ and/or PQS could serve as a signal(s) that informs phage JBD44^39*^ to lysogenize its host bacterium, rather than lyse the host. We say this because the transition from the initial lysis period to the later lysogeny period correlates with production of these compounds. Phages also encode their own QS systems that promote the transition from lysis to lysogeny when susceptible hosts become scarce (18). Thus, QS clearly shapes the outcomes of phage-host interactions. Because QS also regulates bacterial pathogenicity, future efforts to investigate phage-host QS interactions during bacterial infections will likely reveal additional cross-domain relationships.

Twenty-two percent of *P. aeruginosa* clinical isolates from cystic fibrosis patients harbor mutations in *lasR* (46). Our findings suggest that the use of phage therapies in this medical context may activate *P. aeruginosa* LasI/R and the RhlI/R and PQS QS signaling systems and thereby render bacteria that are not immediately killed by phage more virulent and, moreover, more prone to resist phage killing via upregulation of the CRISPR-*cas* defense system. Thus, efforts to simultaneously inhibit LasI/R, RhlI/R, and PQS signaling may be required to increase the efficacy of phage therapies and reduce the ability of the bacteria to deploy anti-phage defenses.

## Materials and Methods

### Bacterial strains, plasmids, and phages

PA14 and mutants were grown at 37 °C with shaking in Luria-Bertani (LB) broth or on LB agar plates solidified with 15g agar/L. The strains and plasmids used in this study are listed in Table S1. To construct chromosomal mutations in PA14 and the PA14 Δ*lasI* strain, DNA fragments flanking the gene(s) to be deleted were amplified by PCR, sewn together by overlap extension PCR, and cloned into pEXG2 (a generous gift from Joseph Mougous, University of Washington, Seattle, WA) using appropriate restriction sites (47). The resulting plasmids were used to transform *Escherichia coli* SM10λ*pir* and were subsequently mobilized into PA14 or the PA14 Δ*lasI* strain via mating. Exconjugants were selected on LB medium containing gentamicin (30 μg/mL) and irgasan (100 μg/mL), followed by recovery of mutants on M9 medium containing 5% (wt/vol) sucrose. Candidate mutants were confirmed by PCR and sequencing. The primers used are listed in Table S2.

### Plaque assay

25 μL of an overnight culture of PA14 or a mutant strain was combined with 5 mL top LB agar (0.8 % agar and 10 mM MgSO_4_ at 50 °C) and plated on LB solidified with 15 g agar/L. Phage lysates were serially diluted in SM buffer (100 mM NaCl, 8 mM MgSO_4_, 50 mM Tris HCl pH 7.5, 0.01% gelatin) and spotted on bacterial lawns. Plates were incubated at 37 °C. In Fig. 1D, the top and bottom agar were supplemented to a final concentration of 2 μM 3OC12-HSL. In Fig. 2B, concentrated 3OC12-HSL (5 μL of 100 μM 3OC12-HSL) was applied to the center of the top agar layer to achieve a high local concentration that did not diffuse into the entire plate. DMSO solvent was likewise added as the control in both cases.

### Isolation of the phage JBD44^39*^ mutant

A spontaneous mutant of phage JBD44 that produced a small clear plaque morphology was isolated by infecting a culture of PA14 Δ*CRISPR* Δ*cas* with phage JBD44 and sub-culturing at a 1:1,000 dilution each day for three days. The cell-free culture fluid from the overnight culture obtained from the third passage was sterilized by vortex with 1% chloroform. This preparation was combined with 25 μL of an overnight culture of PA14 Δ*CRISPR* Δ*cas* and 5 mL of soft LB agar, overlaid on LB agar plates, and incubated at 37 °C overnight. Phage JBD44 forms large turbid plaques with halos on lawns of PA14. Plaques were screened for those that were clear, indicating a spontaneous mutation favoring lysis had occurred. Plaques harboring putative mutant JBD44 phages were serially streaked three times on lawns of the PA14 Δ*CRISPR* Δ*cas* strain and single plaques were isolated. The mutant phage JBD44 from these isolates was sequenced using MiSeq and analyzed relative to the phage JBD44 reference sequence NC_030929.1. A point mutation in *gp39* was identified, thus the mutant was named JBD44^39*^.

### Determination of the phage JBD44^39*^ integration site

DNA was isolated from the PA14 Δ*lasI* strain lysogenized by phage JBD44^39*^ using the DNeasy Blood & Tissue kit (Qiagen) followed by treatment with 100 μg/mL RNAse A (Qiagen). The DNA was sequenced using MiSeq and analyzed relative to the PA14 reference sequence NC_008463.1.

### Microscopy

The surface profiles of lawns of PA14 and the PA14 Δ*lasI* stain infected with phage JBD44^39*^ that had been grown for 36 h were analyzed using a Leica DCM 3D Micro-optical System. A 10× objective was used with a z step size of 2 μm. A three-point flattening procedure was first performed on the agar surface to level the images. All images were generated by the LeicaMap software associated with the instrument.

### qRT-PCR

Bacterial lawns or plaques were harvested and combined with RNAprotect Bacteria Reagent (Qiagen). RNA was purified using NucleoSpin RNA (Macherey-Nagel) and DNase-treated (RNaseOUT, Thermo Fisher). cDNA was synthesized using SuperScript™ IV Reverse Transcriptase with Random Primers (both from Thermofisher) and quantified using PerfeCTa SYBR Green FastMix Low ROX (Quanta Biosciences).

### Liquid chromatography-mass spectrometry detection of HHQ and PQS levels

50 mg of bacterial lawns or plaques were harvested and homogenized in 1 mL LB broth using a piston. The samples were incubated for 10 min at room temperature and subjected to centrifugation for 2 min at 12,000 RPM. The resulting clarified supernatants were sterilized using 10 kDa centrifugal filter units (Amicon Ultra). These preparations were combined 1:1 with MeOH. HHQ and PQS (Sigma) standards were prepared in 50% MeOH. Liquid chromatography-mass spectrometry was performed to quantify the compounds using a Shimadzu HPLC system as described earlier (48).

### Data analysis

Each experiment was performed at least three times. The results are shown as means ± standard deviations. The time-lapse movie was made using ImageJ.

## Acknowledgements

We thank members of the B.L.B. group for helpful suggestions, Dr. Sampriti Mukherjee for sharing strains, Dr. Jing Yan for assistance with Leica imaging, and Dr. Thomas J. Silhavy for input. We thank Dr. Tharan Srikumar of the Princeton University Proteomics and Mass Spectrometry Core Facility for liquid chromatography-mass spectrometry analyses and Dr. Arshnee Moodley and the NGS-MiSeq facility at the University of Copenhagen for sequencing. We are grateful to Dr. George O’Toole, Dr. Joseph Mougous, and Dr. Joseph Bondy-Denomy for providing bacterial strains and phages. This work was supported by the Howard Hughes Medical Institute, NIH Grant 2R37GM065859, and National Science Foundation Grant MCB-1713731 (to B.L.B.); and Lundbeck Foundation grants R220-2016-860 and R264-2017-3936 (to N.M.H-K.). The funders had no role in study design, data collection and interpretation, or the decision to submit the work for publication.

